# Virological traits of the SARS-CoV-2 BA.2.87.1 lineage

**DOI:** 10.1101/2024.02.27.582254

**Authors:** Lu Zhang, Alexandra Dopfer-Jablonka, Inga Nehlmeier, Amy Kempf, Luise Graichen, Noemí Calderón Hampel, Anne Cossmann, Metodi V. Stankov, Gema Morillas Ramos, Sebastian R. Schulz, Hans-Martin Jäck, Georg M. N. Behrens, Stefan Pöhlmann, Markus Hoffmann

## Abstract

The highly mutated SARS-CoV-2 BA.2.87.1 lineage was recently detected in South Africa, but its transmissibility is unknown. Here, we report that BA.2.87.1 efficiently enters human cells but is more sensitive to antibody-mediated neutralization than the currently dominating JN.1 variant. Acquisition of adaptive mutations might thus be needed for high transmissibility.

## Main text

The emergence and rapid global dominance of the highly mutated Omicron variant in 2021 and its sublineage JN.1 (a derivative of BA.2.86) in 2023 reveals that novel, antigenically distinct variants can rapidly reshape the now fading COVID-19 pandemic. At the end of 2023, a novel SARS-CoV-2 lineage, BA.2.87.1, was detected in eight patients in South Africa and one traveler entering the USA. The BA.2.87.1 lineage harbors 65 mutations in the spike (S) protein (relative to the virus that circulated in Wuhan in early 2020 (Figure 1A)), which facilitates viral entry into cells and constitutes the key target for neutralizing antibodies. However, it is unknown whether these mutations are compatible with robust entry into human cells and allow for efficient antibody evasion. We addressed these questions using pseudovirus particles (_pp_) bearing the SARS-CoV-2 S protein, which adequately model key aspects of SARS-CoV-2 entry into host cells and antibody-mediated neutralization (1). Besides particles bearing BA.2.87.1 S (BA.2.87.1_pp_), we included particles pseudotyped with the S proteins of the B.1 lineage (B.1_pp_), which circulated early in the pandemic, the XBB.1.5 lineage (XBB.1.5_pp_), which served as the target lineage for adaptation of the latest COVID-19 mRNA vaccines (2), and the currently prevailing JN.1 lineage (JN.1_pp_). All S proteins were efficiently processed and incorporated into particles (Figure 1B).

**Figure 1:**
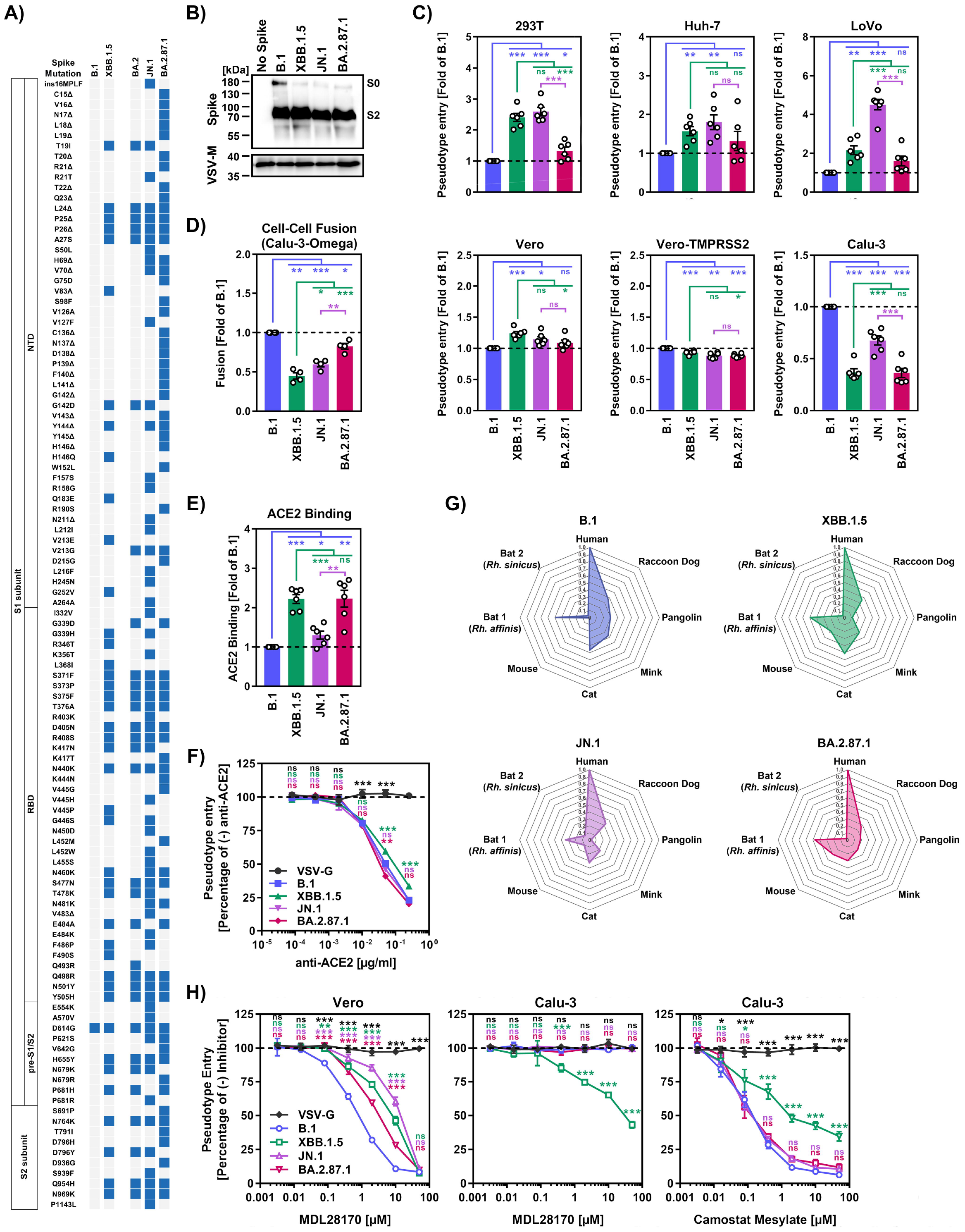
Host cell entry properties of the SARS-CoV-2 BA.2.87.1 lineage. **(A)** S protein mutations of B.1, XBB.1.5, BA.2, JN.1 and BA.2.87.1 compared to the Wuhan-Hu-01 isolate. Abbreviations: NTD, N-terminal domain; RBD, receptor-binding domain, pre-S1/S2, region between RBD and S1/S2 cleavage site. **(B)** Processing and particle incorporation of the BA.2.87.1 S protein. Presented are representative data from a single biological replicate and results were confirmed in five additional biological replicates. **(C)** Entry efficiency of the BA.2.87.1 lineage. Pseudotype particles harboring the indicated S proteins were inoculated onto the indicated cell lines and entry was analyzed. Presented are mean data from six biological replicates, conducted with four technical replicates, with cell entry normalized against particles harboring the B.1 S protein (set as 1). Error bars represent the standard error of the mean (SEM). **(D)** Cell-cell fusion capacity of the BA.2.87.1 lineage. Presented are the mean data from four biological replicates, conducted with three technical replicates. Fusion driven by the B.1 S protein was set as 1. Error bars indicate the SEM. **(E)** Soluble human ACE2 binding by the BA.2.87.1 S protein. Presented are mean ACE2 binding data from six biological replicates, conducted with a single technical replicate, and ACE2 binding was corrected for S protein cell surface expression and normalized using the B.1 S protein as reference (= 1). Error bars indicate the SEM. **(F)** Impact of ACE2 blockade on cell entry of the BA.2.87.1 lineage. Pseudotype particles harboring the indicated S proteins were inoculated onto Vero cells that were preincubated with different concentration of an ACE2-blocking antibody and entry was analyzed. Presented are mean data from three biological replicates, conducted with four technical replicates, with cell entry in the absence of antibody used as reference (set as 100%). Error bars represent the SEM. **(G)** Utilization of mammalian ACE2 orthologues by the BA.2.87.1 lineage. Particles bearing the indicated S proteins were inoculated onto BHK-21 cells expressing the indicated ACE2 orthologues following transfection and entry efficiency was analyzed. Net plots present the mean data from three biological replicates, conducted with four technical replicates, and data were normalized to human ACE2 (set as 1). **(H)** Dependency of BA.2.87.1 lung cell entry on TMPRSS2. Pseudotype particles harboring the indicated S proteins were inoculated onto Vero and Calu-3 cells that were preincubated with MDL28170 or camostat mesylate and entry was analyzed. Presented are mean data from three biological replicates, conducted with four technical replicates, with cell entry in the absence of inhibitor used as reference (set as 100%). Error bars represent the SEM. For panels C, D and E statistical significance was analyzed by two-tailed students’ t-test with Welch correction, while for panels F and H, statistical significance was analyzed by two-way ANOVA with Dunnett’s posttest (p > 0.05, not significant [ns]; p ≤ 0.05, ^*^; p ≤ 0.01, ^**^; p ≤ 0.001, ^***^)

### BA.2.87.1 efficiently enters and fuses human cells

All particles efficiently entered a panel of six cell lines (Figure 1C). BA.2.87.1_pp_ entered 293T (human, kidney), Huh-7 (human, liver), LoVo (human, colon) and Vero cells (African green monkey, kidney, ± S protein-priming protease TMPRSS2) with similar efficiency as B.1_pp_ and JN.1_pp_, except for 293T and LoVo cells, which were more susceptible to JN.1_pp_. For Calu-3 cells (human, lung), entry of B.1_pp_ was highest, followed by JN.1_pp_, whereas XBB.1.5_pp_ and BA.2.87.1_pp_ entry was less efficient.

The ability of the S protein to fuse infected with uninfected cells is believed to contribute to COVID-19 pathogenesis (4-6), which is why we assessed the capacity of BA.2.87.1 S to drive cell-cell fusion using a split beta-galactosidase reporter assay (Figure 1D and Supplementary figure 1C). BA.2.87.1 S displayed significantly higher cell-cell fusion capacity than XBB.1.5 S and JN.1 S, reaching levels observed for B.1 S protein.

### BA.2.87.1 efficiently utilizes human and animal ACE2 as entry receptors

Next, we analyzed the ability of BA.2.87.1 S to engage the SARS-CoV-2 receptor ACE2 and found that BA.2.87.1 S and XBB.1.5 S bound soluble human ACE2 with comparable efficiency while ACE2 binding of B.1 S and JN.1 S was significantly reduced (Figure 1E). However, antibody-mediated of ACE2 engagement did not reveal major differences in ACE2 dependency for cell entry by the different S proteins (Figure 1F). Moreover, all four S proteins could comparably utilize diverse mammalian ACE2 orthologues as entry receptors, with the exception of pangolin ACE2 (highest for B.1_pp_) and mouse efficiency (lowest for B.1_pp_) (Figure 1G). Thus, the BA.2.87.1 lineage efficiently binds human ACE2 and robustly enters and fuses human cells, although entry into Calu-3 lung cells is reduced compared to JN.1.

### Lung cell entry of BA.2.87.1 depends on TMPRSS2

Most Omicron sublineages show a reduced capacity to employ TMPRSS2 for cell entry, which has been linked to diminished lung cell entry and reduced virulence (8-10). Therefore, we evaluated the dependency of BA.2.87.1_pp_ on TMPRSS2 for lung cell entry using the cathepsin L inhibitor MDL28170 and the TMPRSS2 inhibitor camostat mesylate (Figure 1H). MDL28170 reduced Vero kidney cell entry of all particles analyzed but had no impact on Calu-3 lung cell entry of B.1_pp_, JN.1_pp_ and BA.2.87.1_pp_, while XBB.1.5_pp_ entry into lung Calu-3 cells was diminished. Camostat mesylate inhibited Calu-3 cell entry of all particles, with entry of B.1_pp_, JN.1_pp_ and BA.2.87.1_pp_ being more affected than entry of XBB.1.5_pp_. Finally, neither of the inhibitors reduced entry of control particles bearing the vesicular stomatitis virus glycoprotein (VSV-G). Thus, BA.2.78.1 deviates from other Omicron sublineage in its ability to efficiently employ TMPRSS2 for lung cell entry.

### Few therapeutic monoclonal antibodies neutralize BA.2.87.1

Recombinant monoclonal antibodies (mAb) have been successfully used for COVID-19 therapy but contemporary SARS-CoV-2 lineages developed resistance against most or all of them (11). Using a panel of twelve mAbs that were previously approved for COVID-19 therapy or are currently under development, we found that five of them (Casirivimab, Tixagevimab, Amubarvimab, Regdanvimab and Sotrovimab), displayed neutralizing activity against BA.2.87.1_pp_ and should constitute suitable treatment options (Figure 2A-B and Supplementary figure 2). In comparison, only Sotrovimab was effective against XBB.1.5_pp_and none of the mAbs neutralized JN.1_pp_.

**Figure 2:**
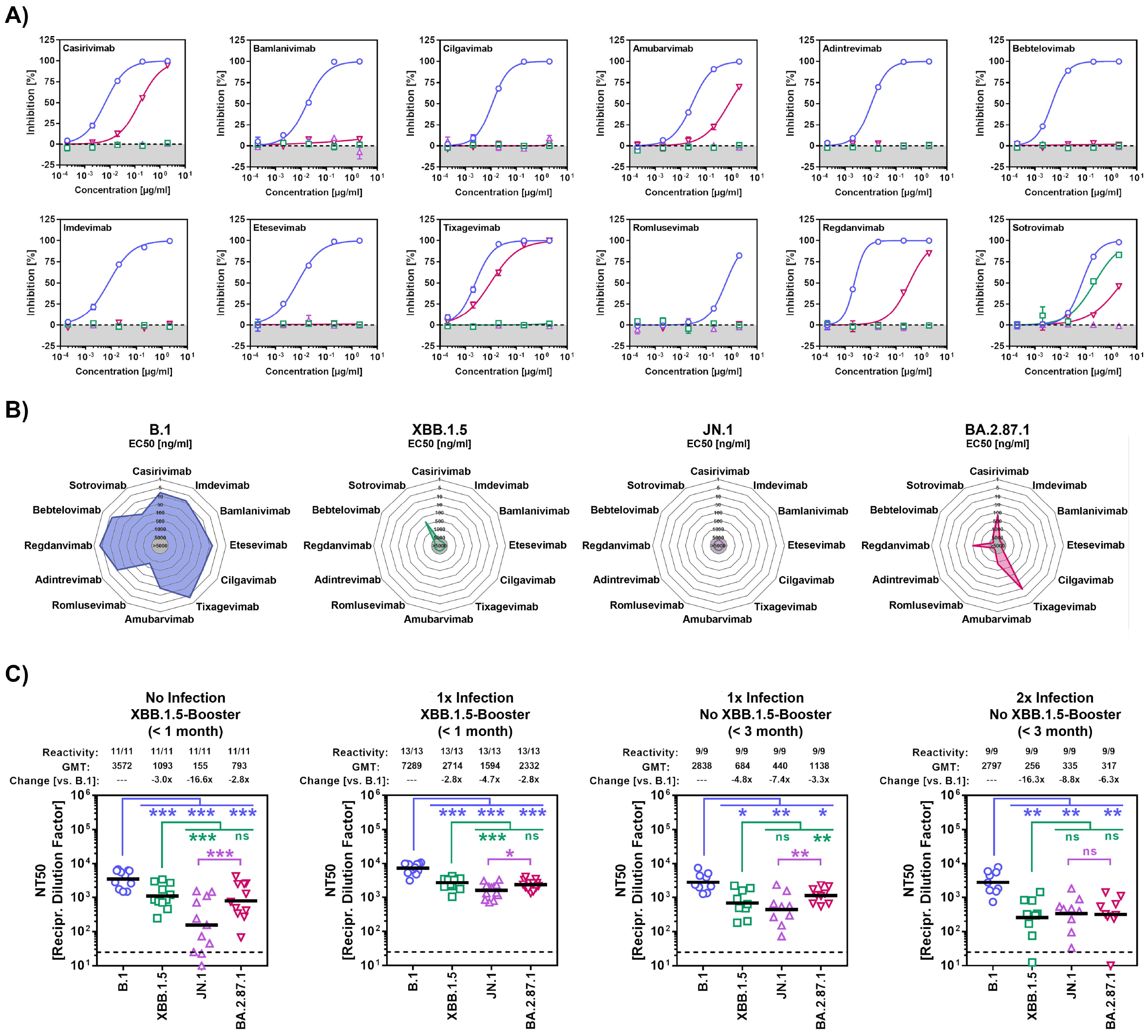
Neutralization sensitivity of the SARS-CoV-2 BA.2.87.1 lineage. **(A)** Sensitivity of the BA.2.87.1 lineage to neutralization by monoclonal antibodies (mAb). Pseudotype particles harboring the indicated S proteins were incubated with different concentrations of the indicated mAbs, before being inoculated onto Vero cells and cell entry was analyzed at 16–18 h post inoculation by measuring firefly luciferase activity in cell lysates. Presented are the mean data from three biological replicates, conducted with four technical replicates, and cell entry was normalized against entry in the absence of mAb (set as 0% inhibition). **(B)** Net plots indicate the effective dose 50 (EC50) values calculated from the data presented in panel A. **(C)** Sensitivity of the JN.1 lineage to neutralization by antibodies in the blood plasma of individuals with different immunization background. Pseudotype particles harboring the indicated S proteins were incubated with different dilutions of plasma, before being inoculated onto Vero cells and cell entry was analyzed at 16–18 h post inoculation by measuring firefly luciferase activity in cell lysates. Cell entry was further normalized against entry in the absence of plasma (set as 0% inhibition) and the neutralizing titer 50 values were calculated based on a nonlinear regression model. Presented are the geometric mean titers (GMT) from a single biological replicate, conducted with four technical replicates. Information above the graphs include response rates (proportion of plasma samples with neutralizing activity), GMT values, and median fold GMT changes compared to particles bearing the B.1 S protein. Please also see Supplementary table 1 and supplementary figures 3 and 4 for additional information. Statistical significance was assessed by Wilcoxon matched-pairs signed rank test (p > 0.05, ns; p ≤ 0.05, ^*^; p ≤ 0.01, ^**^; p ≤ 0.001, ^***^).

### Less neutralization evasion by BA.2.87.1 compared to JN.1

Finally, we studied the sensitivity of BA.2.87.1 to neutralization by antibodies induced upon vaccination or vaccination plus breakthrough infection. Cohorts 1 and 2 included participants who recently received the XBB.1.5-adapted COVID-19 mRNA vaccine from BioNTech (raxtozinameran). Members of cohort 1 had no history of SARS-CoV-2 infection while members of cohort 2 had documented SARS-CoV-2 infection between 01/2022 and 03/2023. Cohorts 3 and 4 included participants without XBB.1.5-booster vaccination, who experienced one (cohort 3) or two (cohort 4) SARS-CoV-2 infections with the most recent infection occurring during the JN.1 wave. Of note, all participants received at least four vaccinations with non-XBB.1.5-adapted COVID-19 vaccines and plasma samples were collected within three months after the last infection or vaccination (Supplementary table 1 and Supplementary figure 3). For all four cohorts, highest neutralizing activity was measured for B.1_pp_ (geometric mean titer = 2797-7289), while neutralization of JN.1_pp_ was lowest (∼5-17-fold reduction compared to B.1_pp_) with the exception of cohort 4 (Figure 2C and Supplementary figure 4). Importantly, although BA.2.87.1_pp_ displayed substantial resistance to antibody-mediated neutralization independent of the cohort analyzed, neutralization evasion was less efficient compared to JN.1_pp_, with the exception of cohort 4.

## Discussion

Our initial virological assessment of the BA.2.87.1 lineage revealed that it efficiently utilizes human and animal ACE2 orthologues as receptors, and robustly enters human cell lines. However, cell entry of BA.2.87.1_pp_ was found to be reduced compared JN.1_pp_. Calu-3 lung cell entry of BA.2.87.1_pp_ was highly dependent on the cellular serine protease TMPRSS2, a trait that is shared with lineages dominating the pre-Omicron era and the recently emerged BA.2.86 lineage (11). With respect to antibody-mediated neutralization, we found that BA.2.87.1 can be neutralized by Casirivimab, Tixagevimab, Amubarvimab, Regdanvimab and Sotrovimab, which could constitute suitable treatment options in case of BA.2.78.1 spread. In addition, BA.2.87.1_pp_ evaded neutralization by antibodies present in the plasma of individuals with diverse immune backgrounds but antibody evasion was reduced compared to JN.1_pp_. Based on the data obtained in this study it seems unlikely that BA.2.87.1 will efficiently spread in regions where JN.1 is dominant. However, BA.2.87.1 may still be able to spread in locations where JN.1 prevalence is low and may acquire additional mutations that improve transmissibility and/or immune evasion.

Limitations of our study include the lack of data for authentic SARS-CoV-2 lineages and small sample sizes of the cohorts, with the latter precluding an analysis of the impact of biological factors (e.g. age, sex, comorbidities, etc.) on neutralization. Nevertheless, this study provides valuable information on the virological traits of the BA.2.87.1 lineage that support political decision makers and medical personnel to determine whether changes in containment and treatment strategies are required.

## Acknowledgements

We gratefully acknowledge the originating laboratories responsible for obtaining the specimens, as well as the submitting laboratories where the genome data were generated and shared via GISAID, on which this research is based. We further thank Roberto Cattaneo, Stephan Ludwig, Andrea Maisner, Stuart G. Turville, and Gert Zimmer for providing reagents. Further, we thank the participants of the CoCo Study for their support and the entire CoCo study team, especially Luis Manthey and Anna Meinecke for help. S.P. acknowledges funding by the EU project UNDINE (grant agreement number 101057100), the COVID-19-Research Network Lower Saxony (COFONI) through funding from the Ministry of Science and Culture of Lower Saxony in Germany (14-76103-184, projects 7FF22, 6FF22, 10FF22) and the German Research Foundation (Deutsche Forschungsgemeinschaft, DFG; PO 716/11-1). L.Z. acknowledges funding by the China Scholarship Council (CSC) (202006270031). A.D.-J. acknowledges funding by the European Social Fund (ZAM5-87006761) and by the Ministry for Science and Culture of Lower Saxony (Niedersächsisches Ministerium für Wissenschaft und Kultur; 14-76103-184, COFONI Network, project 4LZF23). H.-M.J. received funding from BMBF (01KI2043, NaFoUniMedCovid19-COVIM: 01KX2021), Bavarian State Ministry for Science and the Arts and Deutsche Forschungsgemeinschaft (DFG) through the research training groups RTG1660 and TRR130, the Bayerische Forschungsstiftung (Project CORAd) and the Kastner Foundation. G.M.N.B. acknowledges funding by German Center for Infection Research (grant no 80018019238), the European Regional Development Fund Getting AIR (ZW7-85151373), and the Ministry for Science and Culture of Lower Saxony (Niedersächsisches Ministerium für Wissenschaft und Kultur; 14-76103-184, COFONI Network, project 4LZF23. The funding sources had no role in the design and execution of the study, the writing of the manuscript and the decision to submit the manuscript for publication. The authors did not receive payment by a pharmaceutical company or other agency to write the publication. The authors were not precluded from accessing data in the study, and they accept responsibility to submit for publication.

## Ethical statement

Plasma samples were collected as part of the COVID-19 Contact (CoCo) Study (German Clinical Trial Registry, DRKS00021152). The CoCo Study and the analysis performed for this study were approved by the Internal Review Board of Hannover Medical School (institutional review board no. 8973_BO-K_2020, last amendment Sep 2023). All study participants provided written informed consent and received no compensation.

## Data availability

Raw data are available upon request. This study did not generate code. All materials and reagents will be made available upon installment of a material transfer agreement.

## Authors’ contributions

Conceptualization: L.Z and M.H.; Methodology: S.P. and M.H.; Investigation: L.Z., I.N., A.K., L.G., and M.H.; Formal analysis: L.Z., M.H., and M.V.S.; Resources: A.D.-J., N.C.H., A.C., G.M.R., S.R.S, H.-M.J., and G.M.N.B.; Funding acquisition: A.D.-J., H.-M.J., G.M.N.B., and S.P.; Writing – original draft: M.H.; Writing – review & editing: all authors.

