## Supplemental information for "Virological traits of the SARS-CoV-2 BA.2.87.1 lineage"

**Supplementary information**

**Materials and methods**

**Cell culture**

The following cell lines were incubated at 37 °C in a humidified atmosphere containing 5% CO_2_. Vero (African green monkey kidney, female, kidney; CRL-1586, ATCC; RRID:CVCL 0574, kindly provided by Andrea Maisner), 293T (human, female, kidney; ACC-635, DSMZ; RRID:CVCL 0063), Vero cells stably expressing TMPRSS2 (Vero-TMPRSS2; JCRB1819, CellBank Australia; RRID:CVCL_YQ49, kindly provided by Stuart G. Turville) and Huh-7 cells (human, male, liver; JCRB, JCRB0403; RRID: CVCL_0336, kindly provided by Thomas Pietschmann) were cultured using Dulbecco's modified Eagle medium (DMEM, PAN-Biotech), supplemented with 10% fetal bovine serum (FCS, Biochrom), 1% of penicillin (final concentration 100 U/ml) streptomycin (final concentration 0.1 mg/ml of) solution (P/S, PAN-Biotech). LoVo cells (human, male, colon; ACC-350, DSMZ; RRID:CVCL_0399) were cultured using Roswell Park Memorial Institute medium (RPMI, PAN-Biotech), supplemented with 10% FCS and 1% P/S solution, whereas Calu-3 cells (human, male, lung; HTB-55, ATCC; RRID:CVCL_0609, kindly provided by Stephan Ludwig) were cultured using DMEM/F-12 medium (GIBCO), supplemented with 10% FCS, 1% P/S solution, 1x non-essential amino acid solution (from 100x stock, PAN-Biotech) and 1 mM sodium pyruvate (PAN-Biotech). Calu-3 cells stably expressing the beta-galactosidase omega fragment (Calu-3-Omega) were generated by retroviral transduction and selection with puromycin (Invivogen). Calu-3-Omega were further maintained in the same medium as parental Calu-3 cells supplemented with 2 µg/ml of puromycin. All cell lines were regularly tested for the absence of mycoplasma contamination and validated by STR analysis, partial sequencing of the cytochrome c oxidase gene, microscopic examination, and/or according to their growth characteristics. Transfection of 293T cells was performed by calcium phosphate precipitation, while BHK-21 cells were transfected using Lipofectamine 2000 (Thermo Fisher Scientific) according to the manufacturers’ instructions.

**Expression plasmids and sequence analysis**

The expression plasmids pCAGGS-DsRed (1), pCAGGS-VSV-G (2), pCG1-sol-ACE2-Fc (3), pQCXIP_human-ACE2-cMYC (4), pQCXIP_raccoon dog-ACE2-cMYC (4), pQCXIP_pangolin-ACE2-cMYC (4), pQCXIP_mink-ACE2-cMYC (4) pQCXIP_cat-ACE2-cMYC (4), pQCXIP_mouse-ACE2-cMYC (4), pQCXIP_Rhinolophus affinis-ACE2-cMYC (4), pQCXIP_Rhinolophus sinicus-ACE2-cMYC (4), pCG1-SARS-CoV-2 B.1 SΔ18 (codon-optimized, C-terminal truncation of 18 amino acid residues, GISAID Accession ID: EPI_ISL_425259) (5), pCG1-SARS-CoV-2 XBB.1.5 SΔ18 (codon-optimized, C-terminal truncation of 18 amino acid residues, GISAID Accession ID: EPI_ISL_16239158) (6), pQCXIP-beta-galactosidase alpha fragment (7) and pQCXIP-beta-galactosidase omega fragment (7) have been described before, while the expression plasmid for SARS-CoV-2 JN.1 SΔ18 (GISAID Accession ID: EPI_ISL_18530042) was generated by introduction of mutation L455S into plasmid pCG1-SARS-CoV-2 BA.2.86 SΔ18 (codon-optimized, C-terminal truncation of 18 amino acid residues, GISAID Accession ID: EPI_ISL_18114953) (8) via overlap-extension PCR and sequence integrity was confirmed by Sanger sequence using a commercial service (Microsynth SeqLab). The pCG1 expression plasmid was a kind gift from Roberto Cattaneo. Information on SARS-CoV-2 lineages and S protein sequences was collected from the GISAID (Global Initiative on Sharing All Influenza Data) (<https://gisaid.org/>) and CoV-Spectrum (<https://cov-spectrum.org/>) databases.

**Production of pseudovirus particles and cell entry studies**

A previously published protocol was employed to generate vesicular stomatitis virus-based pseudovirus particles bearing SARS-CoV-2 S proteins (9). First, 293T cells transfected to express the respective S protein, vesicular stomatitis virus glycoprotein (VSV-G) or DsRed (negative control) were inoculated with VSV-G-transcomplemented VSV*ΔG(FLuc) (kindly provided by Gert Zimmer) (10). Following an incubation period of 1 h at 37 °C and 5% CO_2_, the supernatant was removed and the cells were washed with phosphate-buffered saline (PBS). Then, medium containing anti-VSV-G antibody (supernatant of I1-hybridoma cells; ATCC no. CRL-2700) was added to all cells except those expressing VSV-G, which instead received medium without antibody. Following an incubation period of 16-18 h, supernatant was transferred into a sterile centrifugal tube and centrifuged for 10 min at 4000 x g in order to pellet cellular debris, before clarified supernatant was used for experiments or stored at ‑80 °C until further use.

Cell entry studies were conducted with target cells seeded in 96-well plates. For experiments assessing the ability of S proteins to use human or animal ACE2 orthologues as receptors, BHK-21 cells were transfected to express the respective ACE2 orthologue (or no ACE2) prior to infection. In case of ACE2-blockade experiments, Vero cells were preincubated (30 min, 37 °C) with different concentrations of anti-ACE2 antibody (Sino Biologicals, 10108-MM36), whereas for experiments addressing the dependency of S protein-driven cell entry on TMPRSS2 and cathepsin L, Vero and Calu-3 cells were preincubated (2 h, 37 °C) with different concentrations of MDL28170 (Santa Cruz) or camostat mesylate (Sigma Aldrich) before pseudovirus particles were added. Following addition of identical volumes of pseudovirus particles, target cells incubated for 16-18 h at 37 °C and 5% CO_2_, before cell entry efficiency was assessed. For this, the activity of virus-encoded firefly luciferase in cell lysates was determined. Cells were lysed by incubated (30 min, room temperature) with PBS containing 0.5% Tergitol (Carl Roth). Then, lysates were transferred to white 96-well plates and luciferase substrate (Beetle-Juice, PJK) was added, before luminescence was quantified using a Hidex Sense plate luminometer (Hidex).

**Analysis of S protein processing and particle incorporation**

Particles bearing the respective S protein (or no S protein, control), were concentrated by high-speed centrifugation (13,300 rpm, 90 min, 4 °C) through a sucrose cushion (20 % w/v sucrose in PBS), before the supernatant was removed and the sucrose cushion was mixed with 1 volume of 2x Sample buffer (0.06 M Tris-HCl, 20% glycerol, 4% SDS, 5% beta-mercaptoethanol, 0.4% bromophenol blue, 2 mM EDTA) and incubated at 96 °C for 15 min. Next, lysates were subjected to SDS-PAGE and proteins were blotted onto nitrocellulose membranes (Hartenstein). The membranes were blocked in PBS-T (PBS with 0.05% Tween-20, Carl-Roth) containing 5% bovine serum albumin (BSA, Carl-Roth) for 30 minutes, before they were probed with primary antibody overnight at 4 °C. For detection of S proteins, anti-SARS-CoV-2 (2019-nCoV) Spike S2 antibody (Biozol, Cat: SIN-40590-T62; rabbit, 1:2000 in PBS-T containing 5% BSA) was used, while the vesicular stomatitis virus matrix protein (VSV-M) was detected as a loading control using an anti-VSV-M [23H12] antibody (Kerafast, Cat: EB0011; mouse, 1:1000 in PBS-T containing 5% skim milk powder). Next, the membranes were washed with PBS-T and incubated for 1 h at 4 °C with horseradish peroxidase-conjugated secondary antibody (S protein detection: anti-rabbit IgG (H+L)-HRPO, Dianova, Cat: 111-035-003; 1:2000 in PBS-T containing 5% skim milk powder; VSV-M detection: anti-mouse IgG (H+L)-HRPO; Dianova, Cat: 115-035-045; 1:2000 in PBS-T containing 5% skim milk powder). Finally, membranes were washed with PBS-T and protein bands were detected using the ChemoCam imaging system with ChemoStar Professional software (Intas Science Imaging Instruments) and an in-house prepared chemiluminescence solution (0.1 M Tris-HCl [pH 8.6], 250 g/ml luminol, 0.1 mg/ml para-hydroxycoumaric acid, 0.3 percent hydrogen peroxide).

**Analysis of S protein-driven cell-cell fusion**

Effector 293T cells transfected to express the respective S proteins (or empty vector) along with the beta-galactosidase alpha fragment were washed, resuspended in medium, seeded on top of Calu-3-Omega (Calu-3 target cells stably expressing the beta-galactosidase omega fragment) and incubated for 18 h. Next, S protein-driven cell-cell fusion was analyzed. For this, a beta-galactosidase substrate (Gal-Screen, Thermo Fisher Scientific) was added and cells were incubated for 90 min, before lysates were transferred into while plates and luminescence was recorded using a Hidex Sense plate luminometer (Hidex).

**Analysis of S protein cell surface expression and ACE2 binding efficiency**

293T cells transfected to express the respective S proteins (or no S protein, control) were washed with PBS, resuspended in PBS-B (PBS with 1% BSA) and pelleted by centrifugation (600 x g, 5 min, room temperature). Next, the supernatant was aspirated and cells were resuspended in PBS-B, before the cell suspension was split into two reaction tubes for the detection of S protein cell surface expression and ACE2 binding.

(i) S protein cell surface expression: Cells were incubated for 1 h at 4 °C with anti-SARS-CoV-2 S2 subunit antibody (Biozol, GTX632604; mouse, 1:100 in PBS-B) in a Rotospin test tube rotator disk (IKA). Thereafter, cells were pelleted by centrifugation (600 x g, 5 min, room temperature) and washed with PBS-B, before they were incubated for 1 h at 4 °C with Alexa Fluor-488-conjugated anti-mouse antibody (Thermo Fisher Scientific, A-10667; 1:200 in PBS-B). Next, cells were pelleted by centrifugation (600 x g, 5 min, room temperature), washed with PBS-B, fixed with 1 % paraformaldehyde solution (30 min, room temperature), washed again and resuspended in PBS-B, before S protein cell surface expression was analyzed using the ID7000 Spectral Cell Analyzer and the ID7000 software (Sony Biotechnology, San Jose, CA, USA).

(ii) ACE2 binding: Cells were incubated for 1 h at 4 °C with soluble human ACE2-Fc (concentrated supernatant of 293T cells transfected with pCG1-sol-ACE2-Fc; 1:20 in PBS-B) in a Rotospin test tube rotator disk (IKA). Thereafter, cells were pelleted by centrifugation (600 x g, 5 min, room temperature) and washed with PBS-B, before they were incubated for 1 h at 4 °C with Alexa Fluor-488-conjugated anti-human antibody (Thermo Fisher Scientific, A-11013; 1:200 in PBS-B). Next, cells were pelleted by centrifugation (600 x g, 5 min, room temperature), washed with PBS-B, fixed with 1 % paraformaldehyde solution (30 min, room temperature), washed again and resuspended in PBS-B, before ACE2 binding was analyzed using the ID7000 Spectral Cell Analyzer and the ID7000 software (Sony Biotechnology, San Jose, CA, USA). Finally, for each S protein ACE2 binding was normalized to S protein surface expression.

**Ethics committee approval and enrolment of study participants**

Collection and analysis of plasma samples was performed as part of the COVID-19 Contact (CoCo) Study (German Clinical Trial Registry, DRKS00021152) and have been approved by the Internal Review Board of Hannover Medical School (institutional review board no. 8973_BO-K_2020, last amendment Sep 2023). All participants provided written informed consent and received no compensation. Of note, the CoCo study is a prospective observational study that monitors anti-SARS-CoV-2 IgG and immune responses in health care professionals at Hannover Medical School and in individuals with potential SARS-CoV-2 contact (11, 12).

**Plasma samples**

*cohort 1*, vaccinated individuals without history of SARS-CoV-2 infection who received the XBB.1.5-adapted booster vaccine as last vaccination (n = 11; age range 25-74 [median = 48 years]; male to female ratio 4:7; sampling 15-21 days post vaccination); *cohort 2*, vaccinated individuals with a history of SARS-CoV-2 infection who received the XBB.1.5-adapted booster vaccine as last vaccination (n = 13; age range 29-62 (median = 44 years); male to female ratio 6:7; sampling 15-17 days post vaccination). *cohort 3*, vaccinated individuals, who did not receive the XBB.1.5-adapted booster vaccine and have a history of one SARS-CoV-2 infection between 11/2023 and 12/2023, (n = 9; age range 31-64 (median = 56 years); male to female ratio 3:6 sampling 44-88 days post infection). *cohort 4*, vaccinated individuals, who did not receive the XBB.1.5-adapted booster vaccine and have a history of two SARS-CoV-2 infections, the last of which occurring between 11/2023 and 12/2023 (n = 9; age range 36-58 (median = 50 years); male to female ratio 2:7 sampling 44-81 days post infection). SARS-CoV-2 S1-specific IgG titers were quantified with the anti-SARS-CoV-2-QuantiVac-ELISA (IgG) (EUROIMMUN) and the SARS-CoV-2 infection-free status of cohort 1 was confirmed by absence of anti-SARS-CoV-2 NCP IgG by the anti-SARS-CoV-2 ELISA (NCP) (EUROIMMUN). Specific information on the plasma samples are summarized in Supplementary table 1. Before experiments, plasma samples were heat-inactivated by incubation at 56 °C for 30 min.

**Neutralization assay**

Neutralization assays were performed based on a published protocol (13). Particles bearing the respective S protein were mixed with different concentrations of mAb (range: 0.2 ng/ml to 2 µg/ml) or dilutions of blood plasma (range: 1:25 to 1:6,400) and incubated for 30 min at 37 °C, before being inoculated onto Vero cells. Following an incubation period of 16-18 h, neutralization efficiency was analyzed. For this, entry was normalized to samples without mAb/plasma (set as 0% inhibition). Further, the mAb concentration or plasma dilution leading to half-maximal inhibition (mAb, Effective concentration 50, EC50; plasma, neutralizing titer 50, NT50) were calculated based on a non-linear regression model. Of note, the thresholds for neutralization-positive mAbs and plasma samples were defined as EC50 ≤ 5 µg/ml (2.5-times the highest mAb concentration tested) and NT50 ≥ 6.25 (25% of lowest plasma dilution tested), respectively.

**Quantification and statistical analysis**

Data were analyzed in Microsoft Excel (part of Microsoft Office Professional Plus, version 2016, Microsoft Corporation) and GraphPad Prism version 8.3.0 (GraphPad Software). Two-tailed Student’s t-test with Welch correction, two-way analysis of variance with Dunnetts’ posttest, and Wilcoxon matched-pairs signed rank test were used to analyze statistical significance (the statistical method of the individual experiments is indicated in the figure legends). Only p values of 0.05 or lower were considered as statistical significant (ns [not significant], p > 0.05; *, p ≤ 0.05; **, p ≤ 0.01; ***, p ≤ 0.001).

**Limitations of the study**

The following limitations apply to our study. First, pseudovirus particles and cell lines were used to assess BA.2.87.1 host cell entry and its neutralization. Thus, our results await formal confirmation with authentic SARS-CoV-2 BA.2.87.1 and primary cell cultures and organoids. Second, the pathogenic potential of BA.2.87.1 remains to be analyzed using *in vivo* models. Third, due to the small sample size for the four cohorts, a detailed analysis on the potential impact of biological factors (e.g. age, gender, or comorbidities) on neutralization efficiency was not possible. Fourth, all plasma samples were collected within three months after the XBB.1.5 booster vaccination or last infection. Thus, it remains to be analyzed whether differences in neutralization efficiencies for the tested SASR-CoV-2 lineages become more or less pronounced after an extended period of time. Fifth, for cohorts 3 and 4 no specific information on the specific SARS-CoV-2 lineages that caused infection is available. Sixth, neutralization sensitivity of the SARS-CoV-2 BA.2.87.1 lineage may differ in cohorts with immune backgrounds distinct from the ones examined in the present study.

**Supplementary tables**

**Table S1: Plasma information**

| **General information** | | | | | | **Vaccination status** | **Infection status** | | |
| --- | --- | --- | --- | --- | --- | --- | --- | --- | --- |
| **Sample ID** | **Cohort** | **Gender** | **Age (years)** | **Days since last immunization^a^** | **IgG (BAU/ml)^b^** | **Vaccination history** | **Infected (Yes/No)** | **Date of infection(s)** | **Most prevalent lineage(s) at the time of infection** |
| 10170 | 1 | Female | 26 | 15 | 6627 | V#1: Yes (n.i.); V#2: Yes (n.i.); V#3: Yes (n.i.); V#4: BNT (B.1/BA.5); V#5: BNT (XBB.1.5) | No | n.a. | n.a. |
| 10179 | 1 | Female | 30 | 16 | 5110 | V#1: BNT (B.1); V#2: BNT (B.1); V#3: BNT (B.1); V#4: BNT (B.1); V#5: BNT (XBB.1.5) | No | n.a. | n.a. |
| 10182 | 1 | Female | 42 | 15 | 2125 | V#1: BNT (B.1); V#2: BNT (B.1); V#3: BNT (B.1); V#4: BNT (B.1/BA.5); V#5: BNT (XBB.1.5) | No | n.a. | n.a. |
| 10186 | 1 | Male | 74 | 15 | 1882 | V#1: AZD; V#2: AZD; V#3: BNT (B.1); V#4: MOD (B.1); V#5: BNT (XBB.1.5) | No | n.a. | n.a. |
| 10192 | 1 | Female | 48 | 16 | 3523 | V#1: BNT (B.1); V#2: BNT (B.1); V#3: BNT (B.1); V#4: BNT (B.1/BA.5); V#5: BNT (XBB.1.5) | No | n.a. | n.a. |
| 10194 | 1 | Male | 61 | 16 | 1114 | V#1: BNT (B.1); V#2: BNT (B.1); V#3: BNT (B.1); V#4: BNT (B.1); V#5: BNT (XBB.1.5) | No | n.a. | n.a. |
| 10197 | 1 | Female | 57 | 16 | 2531 | V#1: BNT (B.1); V#2: BNT (B.1); V#3: BNT (B.1); V#4: BNT (B.1); V#5: BNT (B.1/BA.5); V#6: BNT (XBB.1.5) | No | n.a. | n.a. |
| 10198 | 1 | Female | 64 | 16 | 6560 | V#1: AZD; V#2: AZD; V#3: BNT (B.1); V#4: BNT (B.1); V#5: BNT (XBB.1.5) | No | n.a. | n.a. |
| 10200 | 1 | Male | 57 | 16 | 2106 | V#1: BNT (B.1); V#2: BNT (B.1); V#3: BNT (B.1); V#4: Yes (n.i.); V#5: Yes (n.i.); V#6: BNT (B.1); V#7: Yes (n.i.); V#8: BNT (XBB.1.5) | No | n.a. | n.a. |
| 10215 | 1 | Male | 38 | 16 | 2202 | V#1: BNT (B.1); V#2: BNT (B.1); V#3: BNT (B.1); V#4: BNT (B.1); V#5: BNT (XBB.1.5) | No | n.a. | n.a. |
| 10223 | 1 | Female | 25 | 21 | 4461 | V#1: AZD; V#2: AZD; V#3: BNT (B.1); V#4: BNT (B.1/BA.5); V#5: BNT (XBB.1.5) | No | n.a. | n.a. |
| 10167 | 2 | Male | 44 | 16 | 1552 | V#1: BNT (B.1); V#2: BNT (B.1); V#3: BNT (B.1); V#4: BNT (XBB.1.5) | Yes | 15.04.2022 | BA.2 |
| 10172 | 2 | Male | 50 | 15 | 3066 | V#1: AZD; V#2: BNT (B.1); V#3: BNT (B.1); V#4: BNT (B.1/BA.5); V#5: BNT (XBB.1.5) | Yes | 15.03.2023 | XBB.1.5, CH.1.1, XBB.1.9 |
| 10178 | 2 | Female | 56 | 15 | 3434 | V#1: BNT (B.1); V#2: BNT (B.1); V#3: BNT (B.1); V#4: BNT (B.1); V#5: BNT (XBB.1.5) | Yes | 29.07.2022 | BA.5 |
| 10180 | 2 | Male | 41 | 16 | 2797 | V#1: AZD; V#2: BNT (B.1); V#3: BNT (B.1); V#4: BNT (B.1/BA.5); V#5: BNT (XBB.1.5) | Yes | 14.02.2022 | BA.1, BA.2 |
| 10185 | 2 | Female | 62 | 16 | 1690 | V#1: BNT (B.1); V#2: BNT (B.1); V#3: BNT (B.1); V#4: BNT (B.1); V#5: BNT (XBB.1.5) | Yes | 13.01.2023 | BA.5, CH.1.1, XBB.1.5 |
| 10188 | 2 | Female | 34 | 16 | 1952 | V#1: BNT (B.1); V#2: BNT (B.1); V#3: BNT (B.1); V#4: BNT (B.1); V#5: BNT (XBB.1.5) | Yes | 07.01.2022 | BA.1 |
| 10189 | 2 | Female | 45 | 16 | 2467 | V#1: AZD; V#2: BNT (B.1); V#3: BNT (B.1); V#4: BNT (B.1); V#5: BNT (XBB.1.5) | Yes | 15.07.2022 | BA.5 |
| 10191 | 2 | Female | 29 | 16 | 1395 | V#1: BNT (B.1); V#2: BNT (B.1); V#3: BNT (B.1); V#4: BNT (XBB.1.5) | Yes | 16.02.2022 | BA.1, BA.2 |
| 10201 | 2 | Male | 43 | 16 | 5395 | V#1: BNT (B.1); V#2: BNT (B.1); V#3: MOD (B.1); V#4: BNT (B.1/BA.5); V#5: BNT (XBB.1.5) | Yes | 15.03.2022 | BA.1, BA.2 |
| 10208 | 2 | Female | 42 | 15 | 1261 | V#1: BNT (B.1); V#2: BNT (B.1); V#3: BNT (B.1); V#4: BNT (B.1); V#5: BNT (XBB.1.5) | Yes | 26.10.2022 | BA.5 |
| 10212 | 2 | Female | 31 | 17 | 3446 | V#1: BNT (B.1); V#2: BNT (B.1); V#3: BNT (B.1); V#4: BNT (B.1); V#5: BNT (XBB.1.5) | Yes | 21.05.2022 | BA.2 |
| 10214 | 2 | Male | 58 | 15 | 6090 | V#1: AZD; V#2: BNT (B.1); V#3: BNT (B.1); V#4: BNT (XBB.1.5) | Yes | 15.03.2023 | XBB.1.5, CH.1.1, XBB.1.9 |
| 10224 | 2 | Male | 44 | 16 | 1552 | V#1: BNT (B.1); V#2: BNT (B.1); V#3: BNT (B.1); V#4: BNT (XBB.1.5) | Yes | 13.05.2022 | BA.2 |
| 10396 | 3 | Female | 60 | 53 | 3275 | V#1: MOD; V#2: Yes (n.i.); V#3: Yes (n.i.) | Yes | 17.12.2023 | JN.1 |
| 10410 | 3 | Male | 31 | 44 | 2796 | V#1: BNT (B.1); V#2: BNT (B.1); V#3: BNT (B.1) | Yes | 27.12.2023 | JN.1 |
| 10445 | 3 | Female | 56 | 79 | 4973 | V#1: AZD; V#2: Yes (n.i.); V#3: Yes (n.i.); V#3: BNT (B.1/BA.5) | Yes | 25.11.2023 | JN.1, BA.2.86.1 |
| 10464 | 3 | Female | 58 | 84 | 3032 | V#1: BNT (B.1); V#2: BNT (B.1); V#3: BNT (B.1) | Yes | 20.11.2023 | JN.1, BA.2.86.1 |
| 10503 | 3 | Female | 64 | 47 | 1597 | V#1: AZD; V#2: BNT (B.1); V#3: BNT (B.1); V#4: BNT (B.1/BA.5) | Yes | 28.12.2023 | JN.1 |
| 10518 | 3 | Female | 38 | 46 | 768 | V#1: AZD; V#2: MOD; V#3: MOD | Yes | 29.12.2023 | JN.1 |
| 10543 | 3 | Male | 50 | 60 | 1522 | V#1: BNT (B.1); V#2: Yes (n.i.); V#3: Yes (n.i.) | Yes | 16.12.2023 | JN.1 |
| 10620 | 3 | Male | 58 | 88 | 2074 | V#1: AZD; V#2: BNT (B.1); V#3: BNT (B.1); V#4: BNT (B.1/BA.5) | Yes | 19.11.2023 | JN.1, BA.2.86.1 |
| 10642 | 3 | Female | 56 | 69 | 5541 | V#1: BNT (B.1); V#2: Yes (n.i.); V#3: Yes (n.i.); V#4: BNT (B.1/BA.5) | Yes | 09.12.2023 | JN.1 |
| 10475 | 4 | Female | 58 | 54 | 5937 | V#1: BNT (B.1); V#2: BNT (B.1); V#3: BNT (B.1) | Yes | 12.03.2022 | BA.1, BA.2 |
|  |  |  |  |  |  |  |  | 20.12.2023 | JN.1 |
| 10504 | 4 | Female | 56 | 44-74 | 2226 | V#1: BNT (B.1); V#2: BNT (B.1); V#3: BNT (B.1) | Yes | xx.08.2022 | BA.5, BE.1.1 |
|  |  |  |  |  |  |  |  | xx.12.2023 | JN.1 |
| 10505 | 4 | Male | 54 | 44-74 | 882 | V#1: BNT (B.1); V#2: BNT (B.1); V#3: BNT (B.1) | Yes | xx.12.2022 | BQ.1.1, BF.7 |
|  |  |  |  |  |  |  |  | xx.12.2023 | JN.1 |
| 10514 | 4 | Male | 49 | 81 | 1305 | V#1: BNT (B.1); V#2: Yes (n.i.); V#3: Yes (n.i.); V#4: BNT (B.1/BA.5) | Yes | 15.05.2022 | BA.2 |
|  |  |  |  |  |  |  |  | 24.11.2023 | JN.1, BA.2.86.1 |
| 10533 | 4 | Female | 57 | 78 | 2647 | V#1: AZD; V#2: BNT (B.1); V#3: BNT (B.1) | Yes | 13.09.2022 | BA.5 |
|  |  |  |  |  |  |  |  | 28.11.2023 | JN.1, BA.2.86.1 |
| 10546 | 4 | Female | 42 | 62 | 1519 | V#1: Yes (n.i.); V#2: Yes (n.i.); V#3: Yes (n.i.) | Yes | xx.04.2022 | BA.2 |
|  |  |  |  |  |  |  |  | 14.12.2023 | JN.1 |
| 10562 | 4 | Female | 48 | 45-75 | 1108 | V#1: BNT (B.1); V#2: BNT (B.1); V#3: BNT (B.1) | Yes | 15.07.2022 | BA.5 |
|  |  |  |  |  |  |  |  | xx.12.2023 | JN.1 |
| 10600 | 4 | Female | 36 | 65 | 2118 | V#1: BNT (B.1); V#2: Yes (n.i.); V#3: Yes (n.i.); V#4: BNT (B.1/BA.5) | Yes | 22.12.2022 | BQ.1.1, BF.7 |
|  |  |  |  |  |  |  |  | 12.12.2023 | JN.1 |
| 10653 | 4 | Female | 50 | 47-77 | 924 | V#1: AZD; V#2: BNT (B.1); V#3: BNT (B.1) | Yes | xx.10.2022 | BA.5 |
|  |  |  |  |  |  |  |  | xx.12.2023 | JN.1 |

Cohort 1: No Infection^c^/XBB.1.5 booster; Cohort 2: One infection//XBB.1.5 booster; Cohort 3: One infection/no XBB.1.5 booster; Cohort 4: Two infections/no XBB.1.5 booster. ^a^: For samples without information on the exact date of infection/vaccination a range is provided; ^b^: Anti-SARS-CoV-2 S1 IgG titers were determined against ancestral SARS-CoV-2; ^c^: SARS-CoV-2 infection-free status of cohort 1 was confirmed by ELISA (= anti-NCP IgG-negative). Abbreviations: AZD, AZD1222/Vaxzevria; BNT (B.1), BNT162b2/Comirnaty; BNT (B.1/BA.5), Comirnaty Original / Omicron BA.4-5; BNT (XBB.1.5) Comirnaty XBB.1.5; MOD (B.1), Spikevax; RU, relative units; ID, identifier; IgG, immunoglobulin G; V#, vaccination; n.a., not applicable; n.i., no information available.

**Supplementary figures**

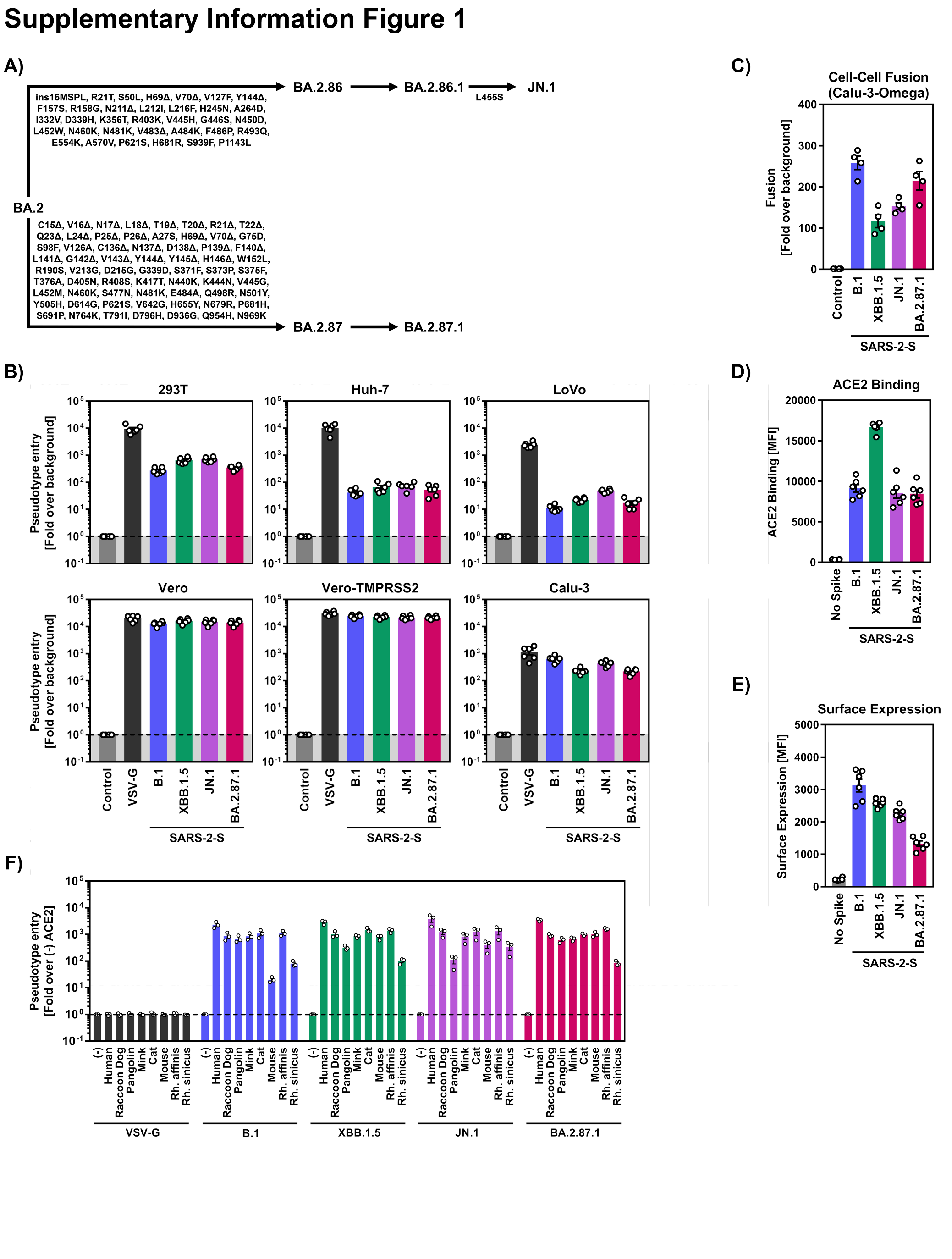

**Figure S1: Host cell entry and ACE2 binding efficiencies of the SARS-CoV-2 BA.2.87.1**

**lineage.**

**(A)** Evolutionary steps leading to JN.1 and BA.2.87.1 (only S protein-specific mutations are shown). **(B)** Cell line tropism and entry efficiency of the SARS-CoV-2 BA.2.87.1 lineage. Particles bearing the indicated S proteins, vesicular stomatitis glycoprotein (VSV-G, positive control), or no viral glycoprotein (negative control) were inoculated onto the indicated cell lines and entry was analyzed at 16–18 h post inoculation by measuring firefly luciferase activity in cell lysates. Presented are the mean data from six biological replicates, conducted with four technical replicates, and cell entry was normalized against the assay background (signals obtained from particles bearing no viral glycoprotein, set as 1). Error bars represent the SEM. **(C)** S protein-driven cell-cell fusion. Effector 293T cells transfected to express the indicated S protein along with the beta-galactosidase alpha fragment were mixed and coincubated with target Calu-3 cells stably expressing the beta-galactosidase omega fragment for 18h. Next, S protein-driven cell-cell fusion was analyzed by quantification of reconstituted beta-galactosidase activity in cell lysates. Presented are the mean data from four biological replicates, conducted with three technical replicates, and fusion was normalized against the assay background (signals obtained after coincubation of target cells with effector cells that did not express S protein, set as 1). Error bars indicate the SEM. **(D)** ACE2 binding efficiency of SARS-CoV-2 S proteins. 293T cells expressing the indicated S proteins following transfection were incubated with soluble human ACE2-Fc and Alexa Fluor-488-conjugated anti-human antibody, before ACE2 binding was analyzed by flow cytometry. Presented are the mean fluorescence intensity (MFI) data from six biological replicates, conducted with a single technical replicate. Error bars indicate the standard deviation (SD). **(E)** Cell surface expression of SARS-CoV-2 S proteins. 293T cells expressing the indicated S proteins following transfection were incubated with anti-SARS-CoV-2 S protein S2 subunit and Alexa Fluor-488-conjugated anti-mouse secondary antibodies, before S protein surface expression was analyzed by flow cytometry. Presented are the mean MFI data from six biological replicates, conducted with a single technical replicate. Error bars indicate the SD. **(F)** Utilization of mammalian ACE2 orthologues by SARS-CoV-2 S proteins. Particles bearing the indicated S proteins or vesicular stomatitis glycoprotein (VSV-G, positive control) were inoculated onto BHK-21 cells expressing the indicated ACE2 orthologues (or no ACE2, control) following transfection and entry efficiency was analyzed at 16–18 h post inoculation by measuring firefly luciferase activity in cell lysates. Presented are the mean data from three biological replicates, conducted with four technical replicates, and changes in cell entry due to ACE2 orthologue expression were calculated using signals obtained from cells expressing no ACE2 as reference (set as 1). Error bars represent the SEM.

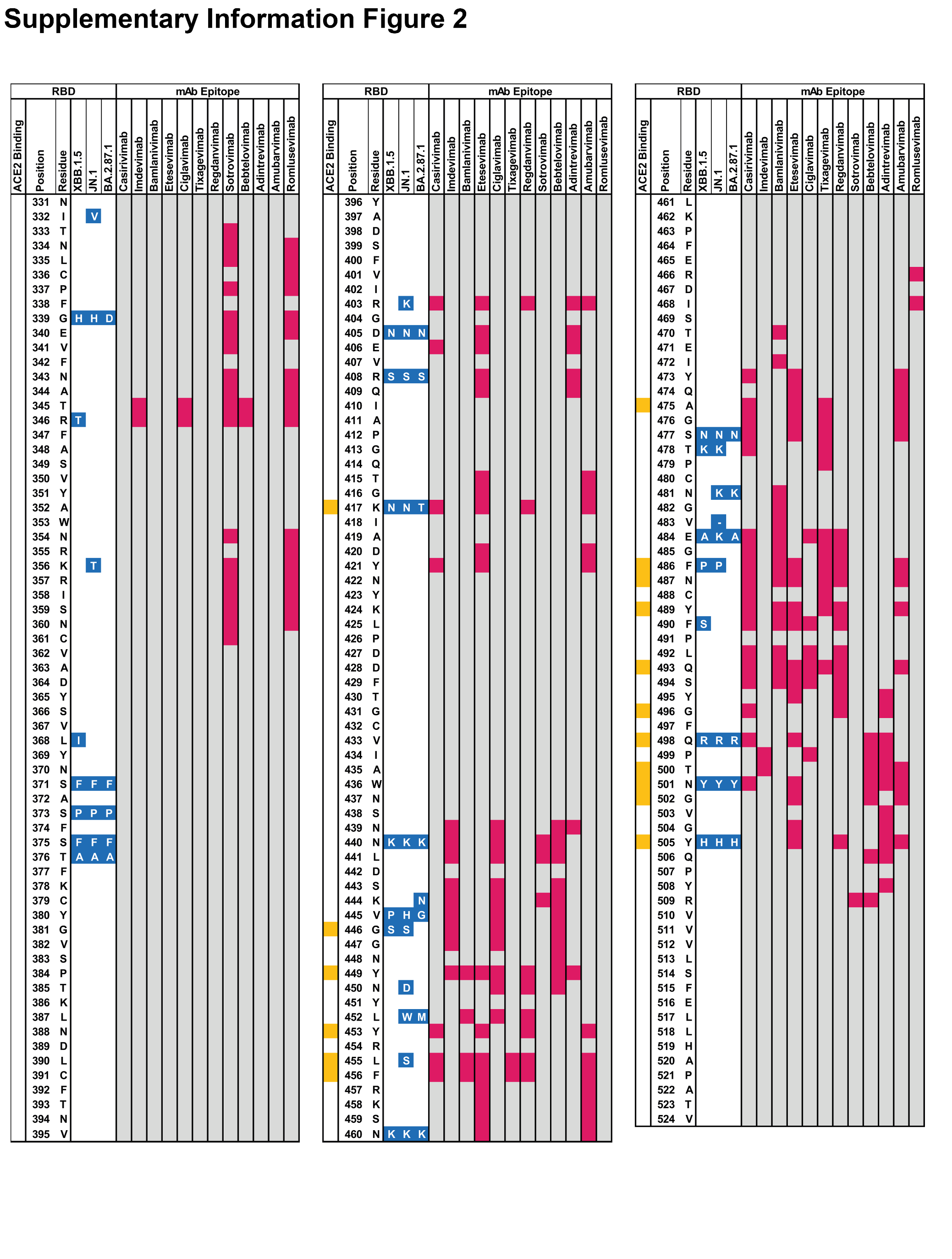

**Figure S2: Overview of BA.2.87.1-specific S protein mutations in the context of epitopes recognized by therapeutic monoclonal antibodies.**

Summary of RBD-specific mutations (blue) in the context of the XBB.1.5, JN.1 and BA.2.87.1 S proteins (numbering according to the S protein of the SARS-CoV-2 Wuhan-Hu-01 isolate). Residues that directly engage ACE2 are highlighted in yellow, while residues that form epitopes recognized by therapeutic monoclonal antibodies are highlighted in pink.

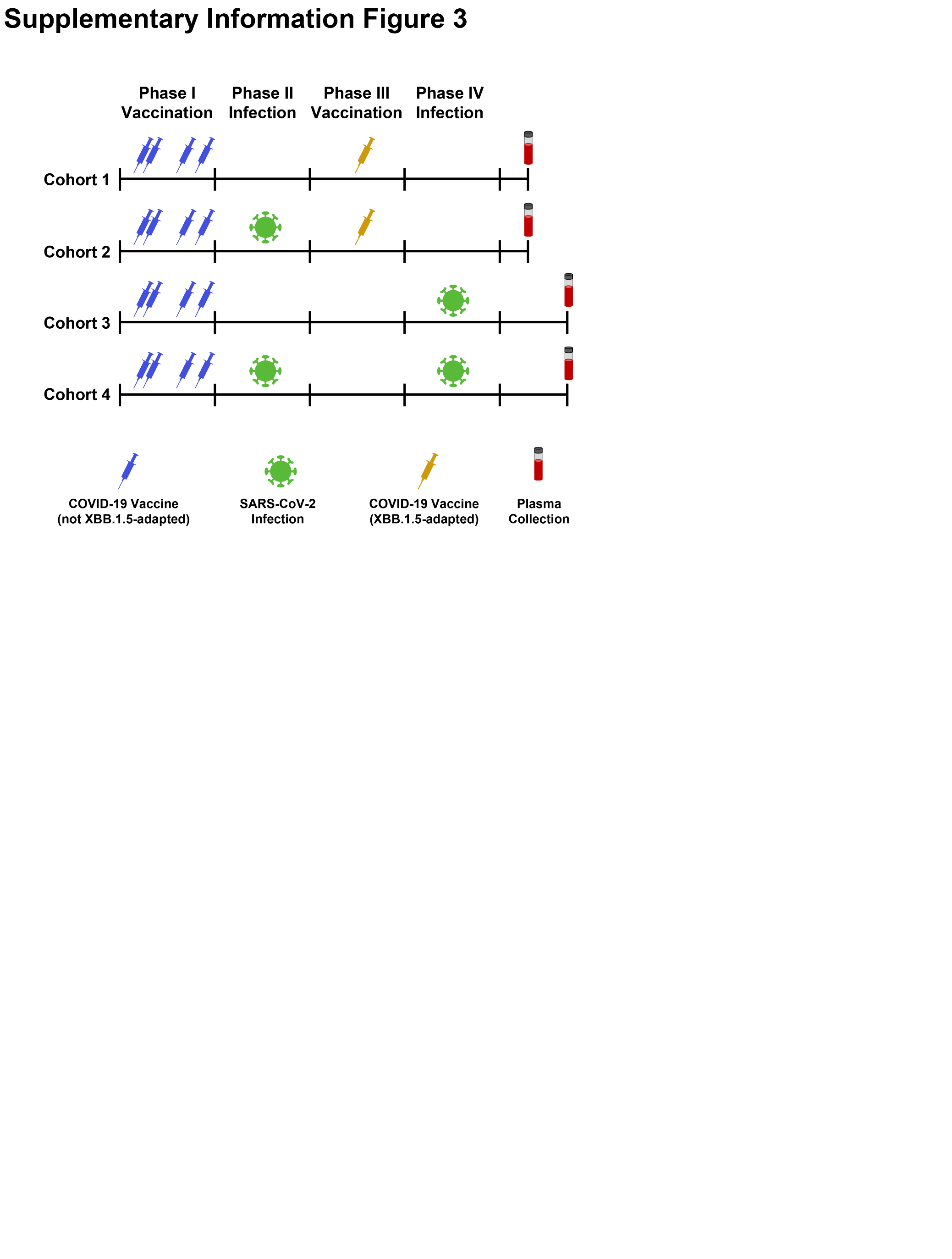

**Figure S3: Immune background of the four cohorts analyzed.**

Cohort 1: Immunization with four to seven doses of non-XBB.1.5-adapted COVID-19 vaccines, followed by a single dose with the XBB.1.5-adapted COVID-19 vaccine of BioNTech/Pfizer. No history of SARS-CoV-2 infection. Cohort 2: Immunization with three to four doses of non-XBB.1.5-adapted COVID-19 vaccines and history of a single SARS-CoV-2 infection between 01/2022 and 03/2023, followed by a single dose with the XBB.1.5-adapted COVID-19 vaccine of BioNTech/Pfizer. Cohort 3: Immunization with three to four doses of non-XBB.1.5-adapted COVID-19 vaccines and history of a single SARS-CoV-2 infection between 11/2023 and 12/2023; no history of vaccination with an XBB.1.5-adapted COVID-19 vaccine. Cohort 4: Immunization with three to four doses of non-XBB.1.5-adapted COVID-19 vaccines and history of two SARS-CoV-2 infections, the last of which happening between 11/2023 and 12/2023; no history of vaccination with an XBB.1.5-adapted COVID-19 vaccine.

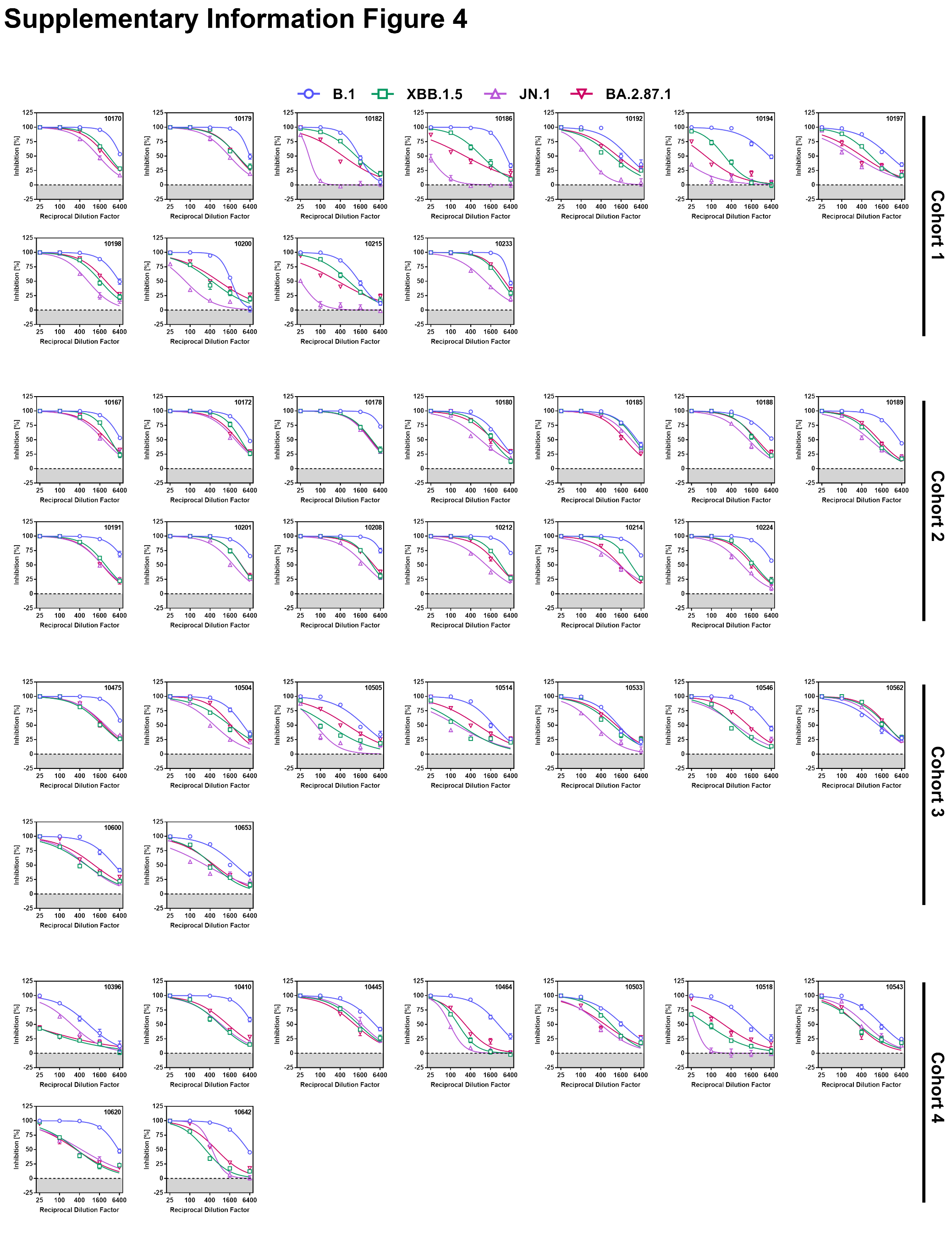

**Figure S4: Neutralization sensitivity of the SARS-CoV-2 BA.2.87.1 lineage.**

Individual neutralization data for plasma samples of the four cohorts. Presented are the mean data from one biological replicates, conducted with four technical replicates, and cell entry was normalized against particles incubated in the absence of plasma (set as 0% inhibition). Error bars indicate the SD.
